# MALER: a web server to build, evaluate, and apply machine learning models for numerical biomedical data analysis

**DOI:** 10.1101/2025.05.09.653008

**Authors:** Zhaoyu Zhai, Zhewei Lin, Qiang Li, Jianbo Pan

## Abstract

The explosive growth of numerical biomedical data poses a challenge in uncovering meaningful insights within from vast omics and clinical data. In recent years, machine learning has emerged as a powerful tool for processing and dissecting numerical biomedical data, making it a popular choice for addressing analytical challenges in biomedical research. Nevertheless, the intricate algorithms and complex optimization processes of machine learning model frameworks have deterred numerous non-machine learning experts. To address the need for comprehensive signatures exploration in biomedical field, we have developed Maler (http://www.inbirg.com/maler/home), a platform designed to facilitate the effortless and efficient application of machine learning models to high-dimensional biomedical data for bioscientists. Maler primarily offers four machine learning modes and a variety of machine learning algorithms to accommodate diverse analysis requirements. Simultaneously, it employs multiple feature selection methods to acquire relevant features and selects the optimal combinations of features based on machine learning models to achieve the best predictive outcomes. Maler is expected to provide a user-friendly and efficient platform for marker discovery, catering to the needs of biologists and clinical experts.

## INTRODUCTION

The continual advancement and innovation in high-throughput sequencing technologies has given rise to a massive influx of omics data, furnishing novel insights and perspectives for research in the field of biomedical science. Recently, extensive analyses of omics data have been conducted to identify specific molecular patterns that serve as biomarkers associated with targeted outcomes of interest, such as clinical diagnosis and disease prognosis (1). Those biomarkers can elucidate the molecular mechanisms of diseases, further promoting precise diagnosis and treatment methodologies. In the contemporary realm of big data, the continuously expanding volume of numerical biomedical data presents significant challenges in terms of in-depth analysis, exploration, and visualization. Machine learning has become pivotal tools to address these challenges (2). By employing techniques such as model training and algorithm optimization, machine learning effectively manages and extracts valuable insights from extensive biomedical datasets. A spectrum of machine learning algorithms, including support vector machines, random forests, and linear regression, has found widespread application in biomarker identification, classification, regression, and survival analysis (3,4). However, the intricacies and complexities involved in the application of machine learning pose significant challenges. The diverse results stemming from various methods, hyperparameters, and data processing choices can be quite bewildering, hindering the integration of machine learning into their research workflows. Additionally, biologists often encounter a deficiency in understanding of machine learning theory and a lack of extensive programming skills. The complexities involved in these processing steps present a notable challenge, making the application of machine learning to their research particularly demanding. Moreover, within the realm of biomedical research, omics datasets typically display highly imbalanced characteristics. The number of features, such as genes and metabolites, often far exceeds the number of samples, leading to the challenge of curse of dimensionality (5). This disparity complicates the accurate learning of patterns within the data, ultimately impacting the performance of the models. Consequently, many researchers combine machine learning methods with feature selection to reduce the challenge of data dimensionality (6). Nevertheless, confronted with the varied and complex landscape of feature selection methods, navigating the challenge of selecting a rapid and efficient selection approach tailored to one’s specific task remains a formidable undertaking. Therefore, the imperative development of user-friendly machine learning analysis pipelines, explicitly designed for the utilization of biologists and clinical researchers, is crucial in overcoming this obstacle and fostering the integration of machine learning into biomedical research.

Currently, there are several open source automated machine learning analysis tools already available, such as Auto-Weka (7), Auto-Sklearn (8) and iLearnPlus (9). These tools integrate various machine learning methods, streamlining the processes of model selection, feature selection and parameter optimization. However, they are not specifically designed to address biological questions, and they may not provide a user-friendly experience for individuals who are not skilled in machine learning. Some of these tools focus solely on hyperparameter optimization. For instance, Auto-Sklearn offers an integrated machine learning pipeline, yet users are still requires to engage in coding for debugging purposes. Conversely, iLearnPlus serves as a comprehensive and user-friendly machine learning automation platform. However, its primary application in nucleic acid and protein sequence analysis limits its utility in disease classification and biomarker discovery.

Considering the complexity of machine learning methods and aiming to bridge the gap in automated omics data analysis, we have developed Maler (Machine Learning Server), a web server that provides a user-friendly and easy-to-use platform for machine learning analysis. Its purpose is to assist biologists in the straightforward and efficient application of machine learning models to high-dimensional biological and medical data. Maler encompasses four major machine learning analysis modules: binary classification, multiclass classification, regression analysis, and survival analysis, addressing diverse user requirements for different data types and analysis needs.

Maler incorporates various common machine learning analysis methods, allowing users to choose the most suitable model tailored to their specific requirements. These modules support a wide range of functionalities, including data normalization, feature selection, machine learning model construction, optimal feature subset selection, model training, performance evaluation and result visualization. Users can obtain diverse analysis results by continually selecting different models and flexibly adjusting parameters. Furthermore, Maler employs comprehensive feature selection methods, systematically constructing various possible feature subsets with combinations of machine learning models. The system can accurately select the optimal machine learning model and further explore potential optimal biomarkers based on multiple performance evaluation results. Maler is meticulously designed to provide users with a comprehensive, efficient, and user-friendly biomedical data automation analysis tool. This empowers researchers to delve deeper into the wealth of information within biomedical data during the era of big data, facilitating a better understanding of potential insights.

## MATERIAL AND METHODS

### Data preprocessing

Data preprocessing is the process of converting raw data into a format suitable for analysis by subsequent machine learning algorithms. The primary objective in this stage is to eliminate noise and interference, thereby improving the overall quality of the data for optimal downstream processing. In Maler, we employ data normalization for uniform preprocessing of input data. Data normalization adjusts feature values to a specified range, enabling the analysis of features with different dimensions within the same dataset. In Maler, we provide three widely utilized standardization methods (Table 1) : Z-score normalization, MinMax normalization, and MaxAbs normalization. In Z-score normalization, features are rescaled to a normal distribution with a mean of 0 and a standard deviation of 1. With MinMax normalization, features are scaled to a unit range between 0 and 1. MaxAbs normalization involves normalizing features based on the absolute value of the maximum, scaling them to a unit range between −1 and 1.

**Table 1:**
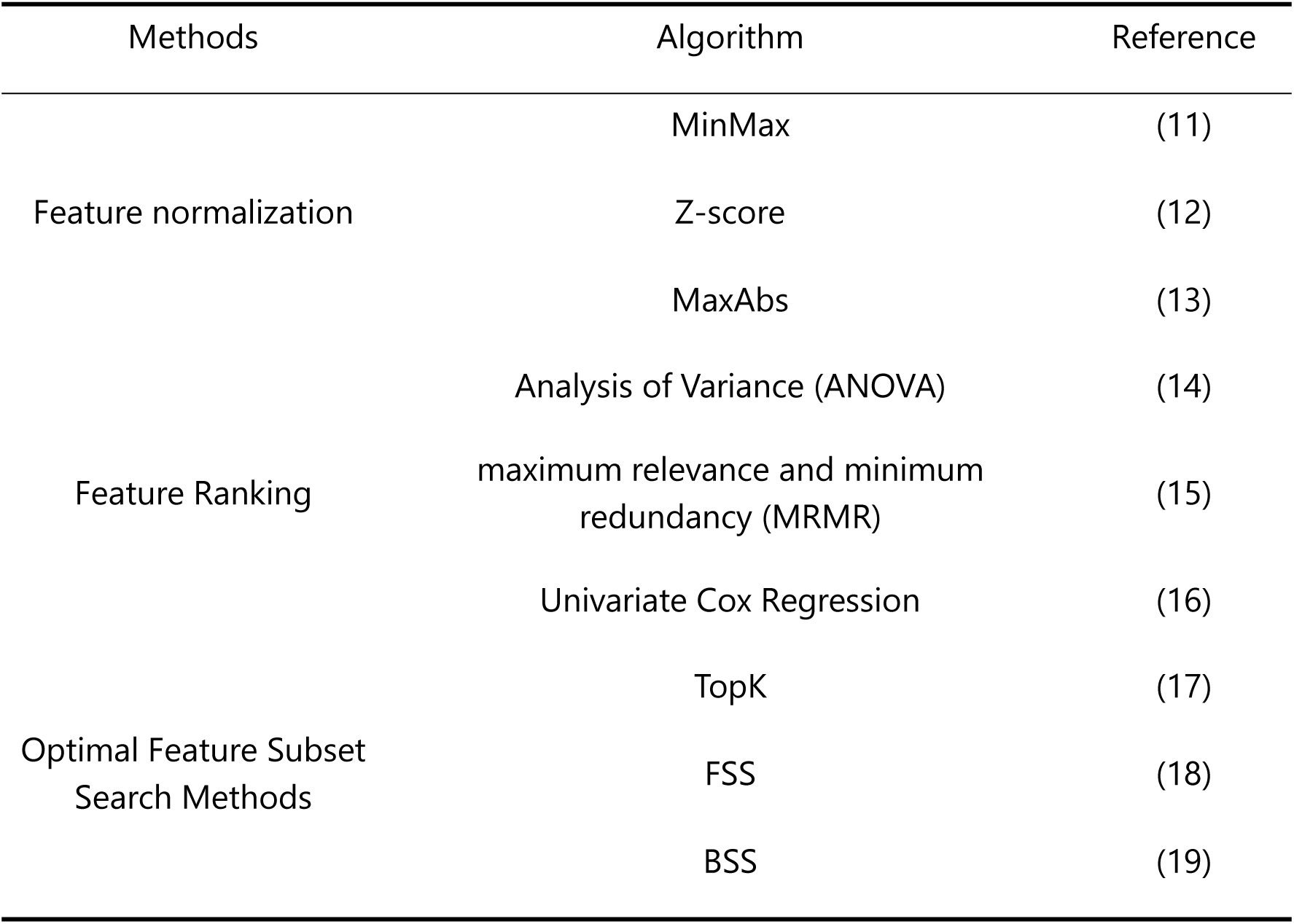
The feature normalization and feature analysis methods provided in Maler.

### Feature analysis

Feature selection is one of the most crucial steps of machine learning, aimed at removing irrelevant, noisy, or redundant features from the original feature set. Its primary goal is to reduce the number of features while preserving the most informative ones, leading to a decrease in computational complexity and an improvement in the predictive performance (10). Maler provides a diverse set of options for feature selection, including two feature ranking methods and three optimal feature subset search methods (Table 1).

Filter and wrapper methods are two commonly used feature selection methods (20,21). Filter methods, commonly relying on the correlation between features, typically employ simple yet efficient approaches to achieve optimal feature selection. The advantage of filter methods lies in their applicability to high-dimensional datasets, with calculations being straightforward and quick (22). A common disadvantage, however, is that the interaction with the classifier and the dependence among features are ignored, which leads to varied classification performance when the selected features are applied to different classification algorithms (23). Wrapper methods integrate feature selection with machine learning models, determining the inclusion or exclusion of features based on their impact on model performance, typically measured by accuracy. While wrapper methods consider the interdependency between features and exhibit a synergistic effect between feature subset and model, they are characterized by computational complexity and a longer processing time (24).

Efficiently and accurately achieving the discovery of optimal features remains a major challenge in machine learning. To strike a balance between efficiency and accuracy, Maler employs a combination of filter and wrapper methods for feature selection. In Maler, we initially employ a filter method to screen features, ranking them based on their correlation with labels, and retaining only the top fifty features (15). This approach allows for the rapid removal of redundant features. Subsequently, we utilize a wrapper method to select the optimal feature subset, seeking the combination within the filtered fifty features that achieves the optimal model performance. This strategy enables the system to conduct feature selection more precisely within the vast feature space, effectively addressing the complexity of biomedical data (25). In classification and regression tasks, Maler offers two feature ranking methods: Analysis of Variance (ANOVA) and the Maximum Relevance and Minimum Redundancy (MRMR). ANOVA ranks features based on the F-statistic, tending to select features with low intra-group variance and high inter-group variance. A higher F-statistic value indicates a stronger correlation between the feature and the target variable (26,27).

The MRMR method aims to identify features with the maximum correlation to the target labels while maintaining minimal redundancy among the selected features (28). For continuous features, it uses the F-statistic to calculate the correlation between features and labels and employs the Pearson correlation coefficient to measure redundancy between features (29). Finally, it ranks the features based on the minimum redundancy maximum relevance criterion derived from mutual information (30). For survival analysis, Maler offers Univariate Cox Regression, ranking features based on the score of the test statistic.

Following the filtering operation, Maler offers three methods: TopK, FSS (31), and BSS for further refining model performance by selecting the optimal subset from the top fifty ranked features.

The TopK method represents a swift feature subset search approach based on prior feature rankings. It involves training a subset model that includes the top k features, with the value of k varying from 1 to 20.

Forward Stepwise Selection (FSS), also known as Sequential Forward Selection, is designed to capture features most relevant to the target variable through a process of iterative evaluation. FSS initiates the search with an empty variable subset, gradually adding variables to a model one at a time. At each step, all the features have not yet been selected are considered for selection, and their impacts on the evaluation score are recorded. The feature with the highest score at each step is then incorporated into the subset. Then, a new step is started, considering the remaining variables. This process continues iteratively, with each subsequent step considering the remaining variables until the predetermined number of features is reached. Additionally, every added feature undergoes a rigorous 10-fold cross validation for model evaluation, ensuring the robustness of the selected features across multiple data subsets.

Backward Stepwise Selection (BSS) method resembles the FSS method, but instead of adding features step by step, it gradually eliminates features from the feature set. The method initiates with a set containing all features. During each iteration, one feature is removed. After each removal, the evaluation score is computed via10-fold cross-validation. Eventually, the feature causing the lowest model evaluation score is removed from the feature set. This elimination process continues until the model comprises only one variable or satisfies the stopping criteria.

### Machine learning model construction

Maler supports online analysis for four distinct target tasks, including binary classification, multiclass classification, regression, and survival analysis. For each task, it provides a respective set of 10, 10, 9, and 5 traditional machine learning algorithms (Table 2), offering users a more comprehensive platform for machine learning analysis to flexibly meet the diverse requirements of different tasks(32). In the process of constructing the mentioned machine learning models, we employed four popular third-party Python packages: scikit-learn (33), XGBoost (34), LightGBM (35), and scikit-survival (36). Specifically, the LightGBM and XGBoost algorithms were implemented using the LightGBM and XGBoost packages, survival analysis-related model algorithms utilized the scikit-survival package, and the scikit-learn library was used for implementing the remaining algorithms.

**Table 2.**
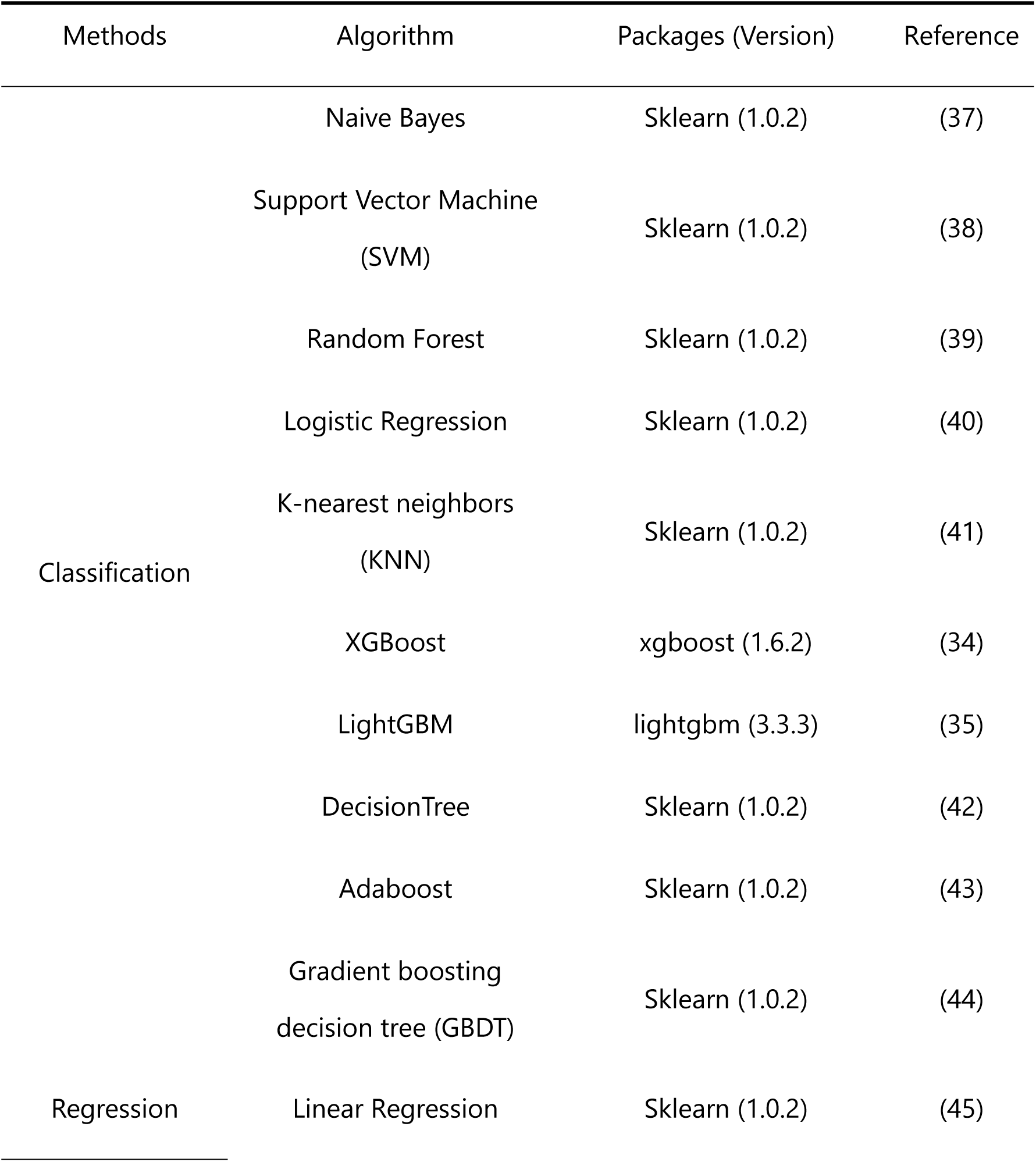
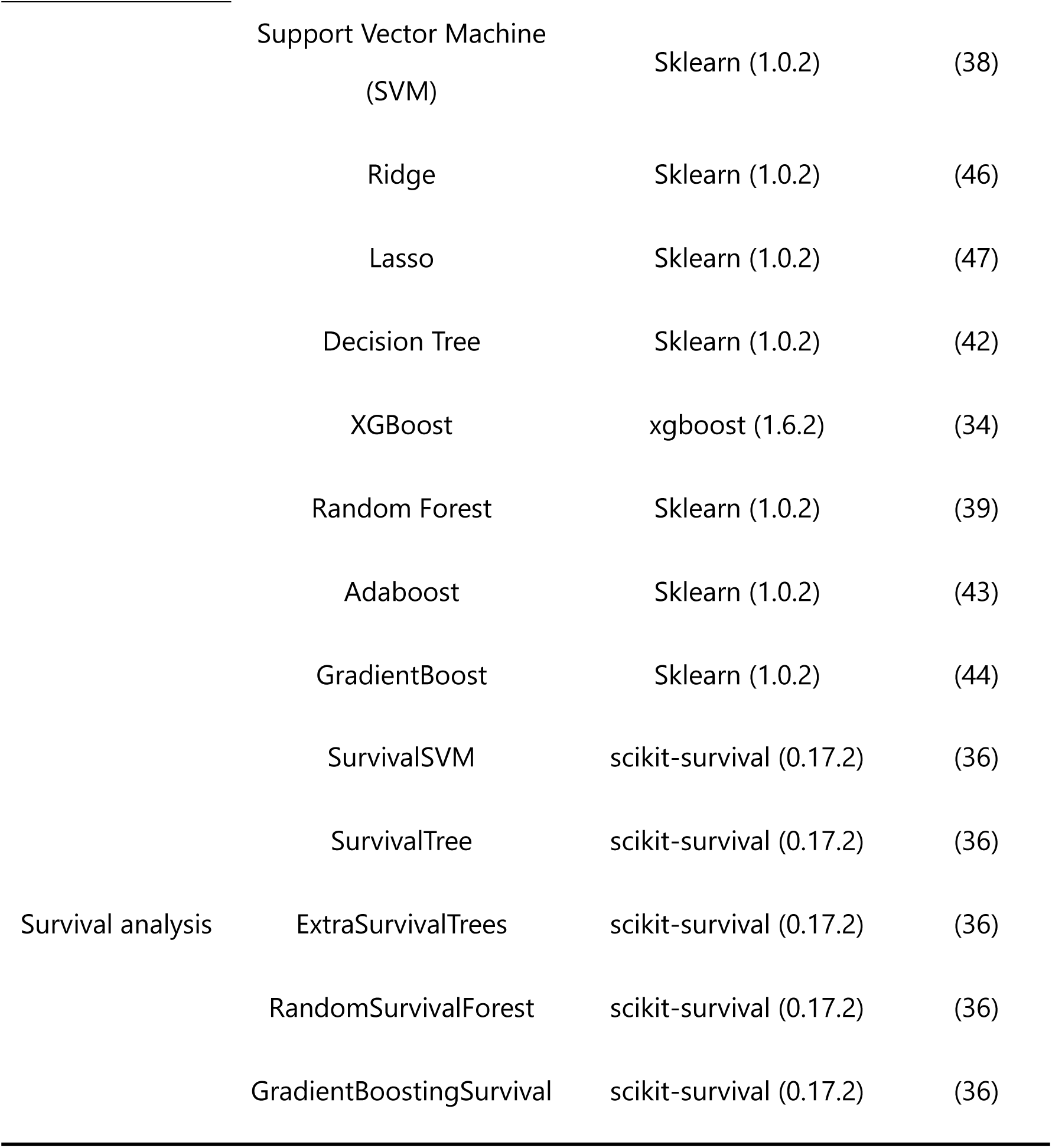
The machine learning algorithms provided in Maler.

Maler offers two machine learning analysis steps: one-click analysis and customized analysis. Through one-click analysis, Maler can automatically analyze all available machine learning models with default parameters simultaneously, displaying the analysis results for all constructed models on the results page. This ensures a more convenient and user-friendly experience, particularly for individuals lacking expertise in machine learning. In the customized analysis module, users can select specific models and adjust parameters according to their preferences. They can also select the “Automatic optimization” option, allowing the system to automatically fine-tune these parameters. For automated parameter optimization, Maler employs a grid search strategy. The system systematically explores various parameter combinations within the specified parameter space to find the optimal configuration in terms of performance.

### Performance evaluation

Maler uses 10-fold cross-validation and a test set evaluation to assess the performance of the constructed machine learning models. In 10-fold cross-validation process, the training set is randomly divided into 10 equally sized folds, with one fold serving as the validation dataset and the remaining 9 folds used as the training dataset to train the machine learning model and optimize its parameters. This process is repeated 5 times to ensure each fold is used as the validation dataset. In each iteration, samples labeled as “training” are utilized for implementing the 10-fold cross-validation test, while samples labeled as “testing” are served as an independent test dataset. This design aimed at evaluating the model’s generalization ability and compare the predictive performance of multiple models (48,49).

For binary classification tasks, we provide five commonly used metrics for quantifying predictive performance. These metrics include Accuracy, Precision, Recall, F1-score, and the Area Under the ROC Curve (AUC), defined as:

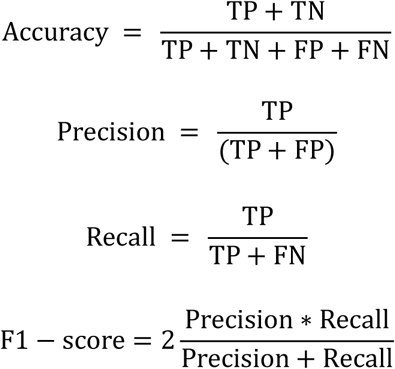

Where TP, FP, TN, and FN represent the numbers of true positives, false positives, true negatives, and false negatives, respectively. The AUC values are calculated based on the receiver-operating-characteristic (ROC) curve, ranging between 0 and 1. Higher

AUC values indicate better predictive performance of the model.

For regression tasks, we provide three commonly used metrics to quantify predictive performance, including R-square, MAE, and MSE, defined as:

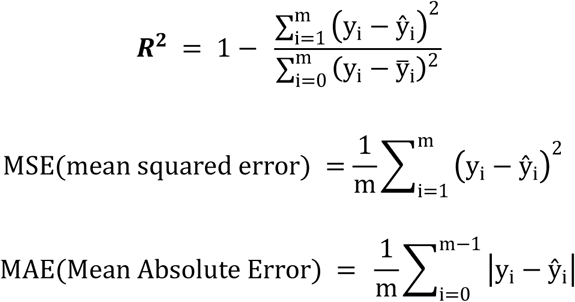

Here, y represents the true label, ŷ represents the predicted label, y-_i_ is the average value, and m denotes the sample size.

For survival analysis tasks, Maler offers the Concordance Index (C-index) (50) as a metric for quantifying predictive performance. The C-index directly reflects the accuracy of the model in ranking survival times. It measures the model’s accuracy in sorting survival times, indicating its ability to correctly predict which sample has a longer survival time among all possible pairs (51). The C-index ranges from 0 to 1, where a higher value indicates better model performance.

### Webserver implementation

Maler is freely accessible at http://www.inbirg.com/maler/home. The online analysis web server framework was implemented using Django (v2.1.8) and deployed on Nginx (v1.14.1) and Gunicorn (v21.2.0) in a Linux environment. The Front-end packages, such as DataTable (v1.10.21), Bootstrap (v4.3.1), Plotly (v2.8.3), and ECharts (v5.0.2), were employed to visually present the results. Python packages, including numpy (v1.21.2), Pandas (v1.3.5), and SciPy (v1.7.3), were employed for performing statistical analyses on the results of predictive model performance.

## RESULTS

Maler covers the four primary steps required for building machine learning models for analysis and prediction: data preprocessing, feature analysis, model construction and performance evaluation and visualization, and new data prediction (Figure 1). We have developed two major modules in Maler to implement these steps: Analysis and Prediction. The Analysis module is primarily used for constructing and training machine learning models, while the Prediction module is focused on applying the trained models to external datasets to validate model performance.

**Figure 1.**
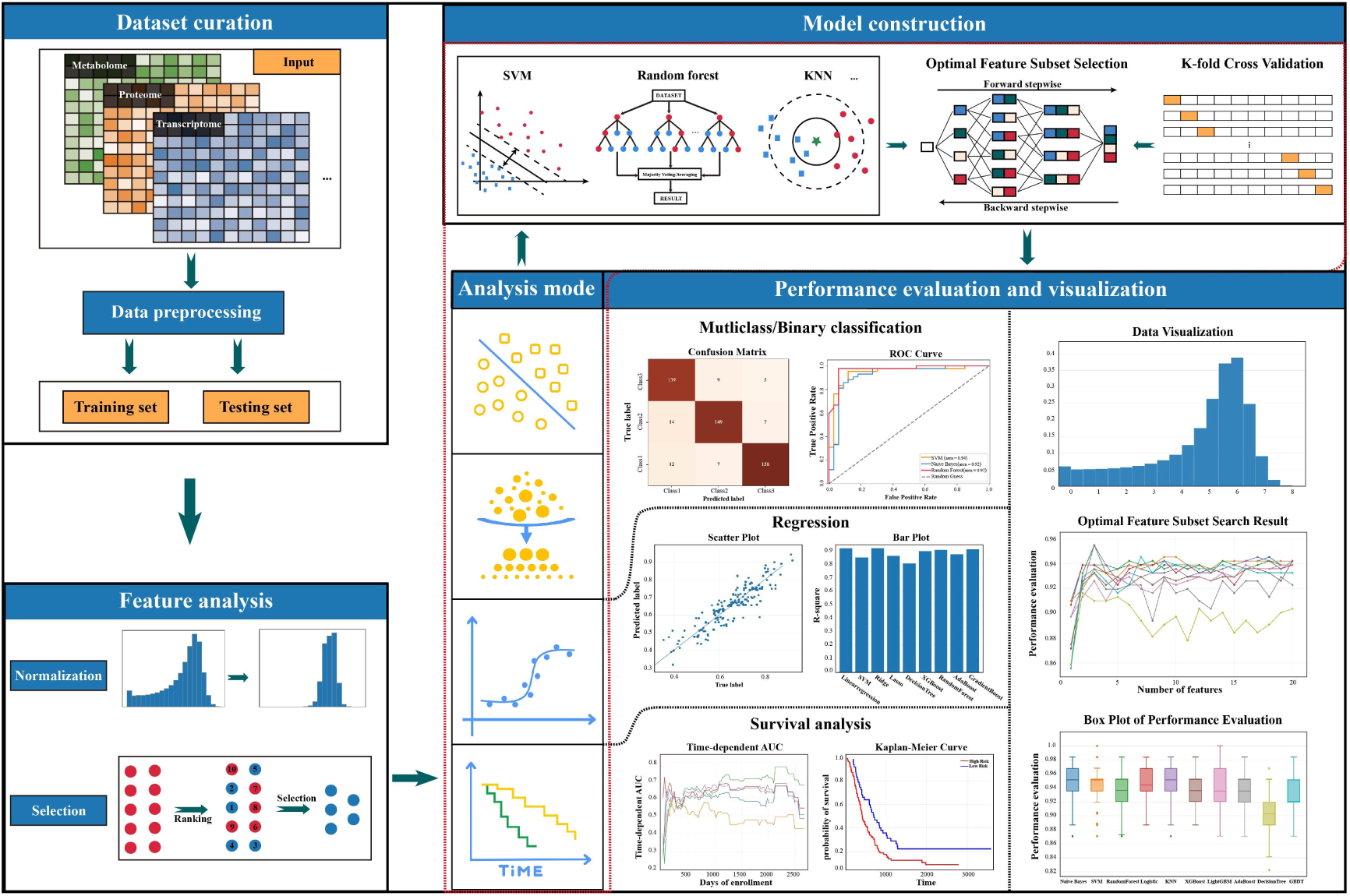
The workflow of the Maler’s machine learning analysis pipeline, including four main built-in modules: Data curation, Feature analysis, Model construction, and Performance evaluation and visualization.

### Data submission

To initiate the analysis process, users first select the analysis mode based on their data features and task objectives, choosing the appropriate machine learning analysis mode for binary classification, multiclass classification, regression, or survival analysis. Subsequently, users select and submit the data file for analysis. Maler supports CSV format data files with feature-by-sample matrix to initiate machine learning analysis. For classification and regression tasks, the first three rows of the input file should respectively provide sample names, sample labels, and the set to which each sample belongs. For survival analysis tasks, the first four rows should provide sample names, status, time, and the set to which each sample belongs. Time represents the follow-up time until the event occurs. Status denotes the patient’s status as “dead” or “alive”.

The data set must contain “training”, and may also contain “testing”. It is essential for users to specify the sample name, sample label (status and time for survival analysis), and features. Sample data of this data type can be downloaded (Figure 2).

**Figure 2.**
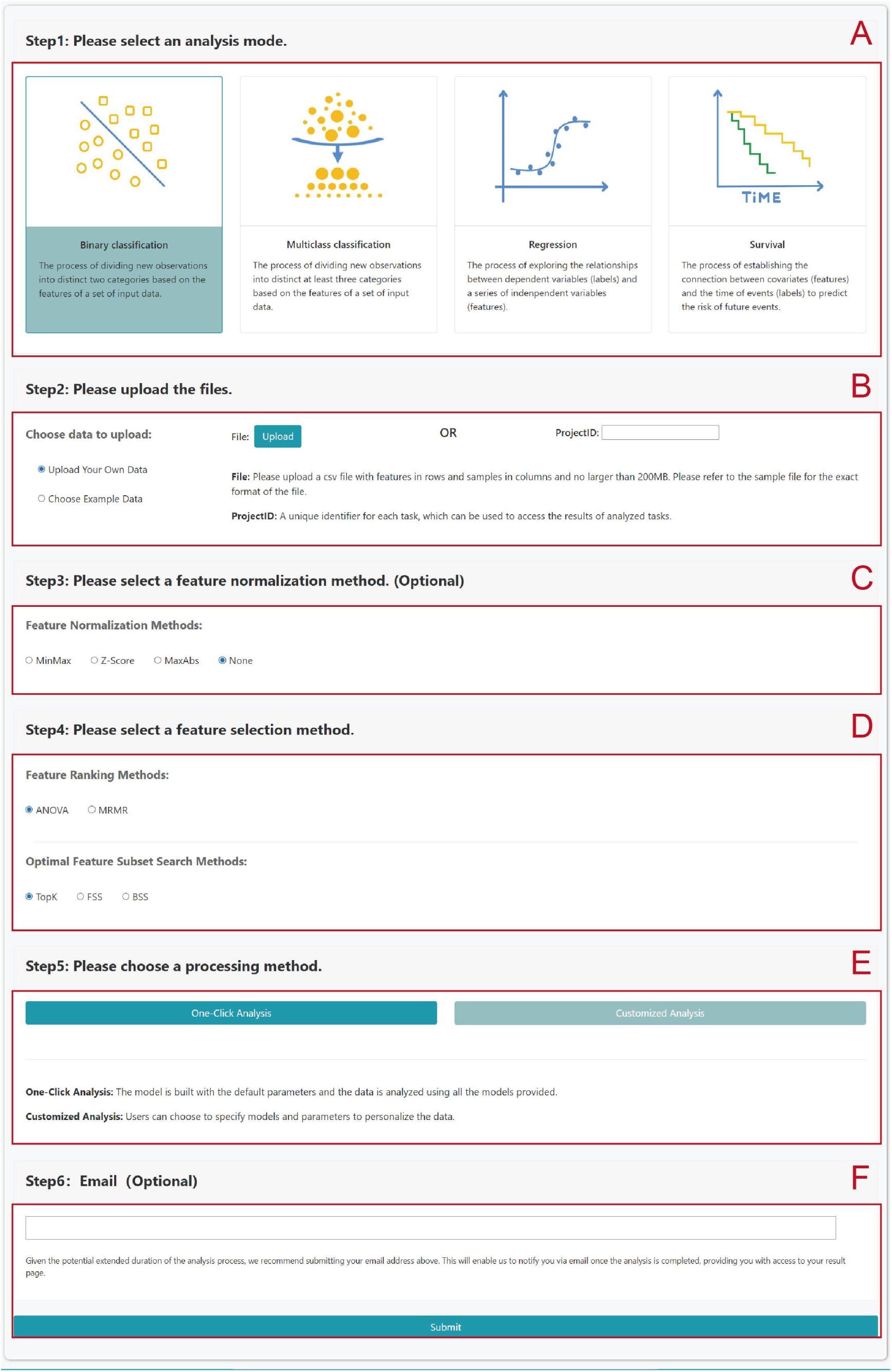
The screenshot shows the analysis workflow of the Maler. Users can generate their desired analysis pipeline in the following six steps: (A) Users can initiate their analysis task by selecting one of four analysis modes. (B) Upload the data to be analyzed. Furthermore, users have the option to submit a projectID, allowing them to compare the performance of different models or parameters using the same dataset. (C-D) The feature analysis process includes feature normalization processing and feature selection methods. (E) Select the machine learning processing method. The one-click analysis method will automatically analyze all available machine learning models simultaneously with default parameters. Alternatively, the customized analysis method allows users to specify the model and parameters for their analysis. (F) To avoid prolonged waiting times on the webpage for analysis results, users can provide an email address. Once the analysis is complete, Maler promptly notifies the user via email and provides a link to the results page.

### Data preprocessing

Following the submission of the data set, the next step involves selecting the data standardization method. Maler offers three data normalization methods to ensure that all features contribute equally with the same numerical scale in subsequent analyses (52). Additionally, users can choose “None” if they prefer not to normalize the data. In cases where the dataset may contain missing values, Maler fills these gaps using the mean from the training set. Finally, the dataset is split into a training set and a test set based on the information provided in the submitted file. If the file doesn’t include information about the sample set, all samples in the data will be assigned to the training set.

### Feature analysis and model construction

After submitting the data, users can select feature ranking methods and optimal feature subset search methods in the “feature selection method” step. Maler offers two feature ranking methods that rank features and select the top 50 features with the highest scores for subsequent model construction. Additionally, Maler offers three optimal feature subset search methods, constructing multiple models by iteratively adding or removing features, and selecting the best results based on evaluation metrics.

In Maler, users benefit from a high level of flexibility, allowing them to specify machine learning models of interest for analysis and dynamically adjust parameters throughout the process. For instance, users can directly specify the penalty and gamma parameters for the radial basis function (RBF) kernel of the SVM classifier. They also have the option to choose automatic parameter optimization or opt for automated tuning within a predefined parameter range. In cases where users possess limited knowledge of machine learning algorithms, Maler offers a one-click analysis feature. Through this functionality, the system constructs and analyzes all available models, empowering users to select the most suitable model based on output results (performance) and refine it as needed. This thoughtful design not only ensures users have complete control over model parameter settings but also provides a convenient automation option. The dual-choice strategy is designed to offer users maximum convenience and flexibility, allowing them to make autonomous decisions based on specific needs and understanding levels.

### Performance evaluation and results visualization

Maler’s computational performance is highly dependent on the dataset size and the selected machine learning algorithm. When users opt to submit a large dataset or train all available machine learning methods, the computation time may extend to several hours. To avoid prolonged waiting times on the web page, users have the option to provide an email address during submission. Once the training of machine learning models is complete, Maler will promptly notify users via email, providing a link to the results page for convenient access.

After completing the aforementioned selection steps and clicking the submit button, Maler processes user data in the backend. Upon completion of preprocessing, the system returns to the preview page, presenting crucial information about the dataset partition (train, test, blind), data distribution, and displaying a tabular representation of some data. Once users confirm the accuracy of the data, they can click the analyze button to initiate the online analysis process for machine learning.

On the results page, Maler presents the results of performance evaluation metrics in a clear table and graphical format, along with showcasing the features selected by the model. This allows users to choose the optimal model based on their specific needs, whether they prioritize achieving the best performance or minimizing the number of features.

Maler employs 5 iterations of 10-fold cross-validation to assess the training set. For all evaluation metrics, the minimum, maximum, average, and variance of the results from 50 runs are calculated and distinctly displayed on the results page. If the user’s dataset includes a test set, Maler further validates the trained model by inputting the test set and computing the corresponding evaluation metrics, thereby providing further confirmation of the model’s performance.

### Data prediction

The models generated by Maler can be exported and saved as model files with the extension “.pkl,” which users can download. In the Predict module, users can upload test data to validate the performance of the trained model on external datasets. When users do not provide label information for the data, predictions are made only for new data, providing corresponding predicted labels for each sample in tabular form. If the dataset includes label information, predictive performance is evaluated, and the corresponding metrics are visualized in chart form on the prediction results page.

Additionally, models saved with traditional machine learning algorithms can be directly applied in the Python environment through the scikit-learn package.

### Case study

Lung cancer is one of the most prevalent types of cancer worldwide and remains a leading cause of cancer-related mortality (53). Non-small cell lung cancers (NSCLCs) approximately account for 85% of lung tumors. NSCLC comprises various subtypes, with the most prevalent being lung adenocarcinoma (LUAD) and lung squamous cell carcinoma (LUSC). These two subtypes exhibit distinct biological behaviors and treatment responses in clinical settings. Consequently, accurate diagnosis of LUAD and LUSC is of paramount significance in tailoring more precise treatment strategies for lung cancer patients (54). Given the variations in histology and gene expression levels between LUAD and LUSC, we showcase the application of Maler in disease classification using expression profile data from LUAD and LUSC as illustrative examples.

In this study, we obtained the RNA-Seq datasets of lung adenocarcinoma (LUAD) and lung squamous cell carcinoma (LUSC) from The Cancer Genome Atlas (TCGA), including 523 LUAD samples and 501 LUSC samples. We transformed the sample expression values into transcripts per million (TPM) and retained only protein-coding genes, resulting in a set of 16,506 original features. The data underwent preprocessing using MinMax normalization, and the ANOVA feature selection method was employed to select the top 50 ranked features, Subsequently, the FSS method was utilized to search for the optimal feature subset. The SVM with radial basis function (RBF) kernel classification algorithm was used for model construction, and the “Automatic optimization” option facilitated automatic parameter tuning in this study.

Finally, after ten-fold cross-validation,the SVM model demonstrated optimal classification performance by selecting 10 features (genes). The average accuracy and AUC values on the training set were distributed as 0.967 and 0.988, respectively, while on the test set, the accuracy and AUC values were 0.964 and 0.985, respectively (Figure 3). In prior research, Chen J. W. et al (55). utilized overlapping feature selection methods to classify and identify biomarkers for LUAD and LUSC, ultimately identifying 17 potential biomarkers with a classification accuracy of 0.929. In comparison to Chen J. W. et al.’s results, Maler achieved superior classification performance using a reduced feature set. Moreover, the entire analysis process in Maler was completed within a mere 10 minutes. Additionally, among the 10 potential biomarkers provided, DSG3, ABCC5, KRT5, ATP1B3, and KRT6C aligned with previous research findings (54,56–58). DAPL1 and SPRR2A were also confirmed to exhibit significant differences in LUAD and LUSC (59), suggesting their potential as biomarkers. This case study highlights the excellence of Maler, showcasing its capacity to generate streamlined yet high-performing models efficiently. As a user-friendly machine learning web server, Maler facilitates profound biomedical data mining and analysis in a straightforward manner.

**Figure 3.**
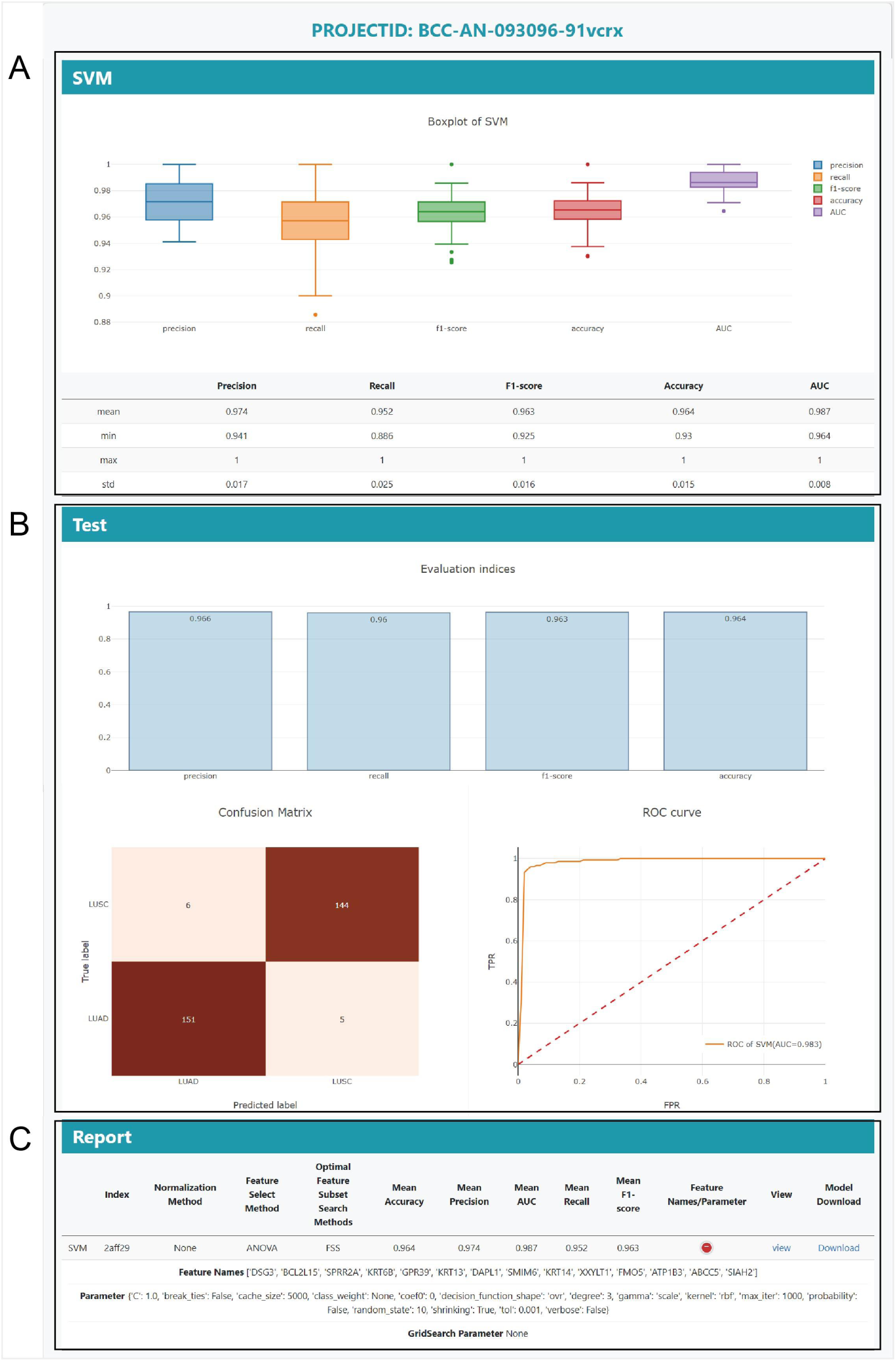
The overview of result page for Lung cancer classification in Maler. (A) presents box plots of classification evaluation metrics after k-fold cross-validation on the training set, revealing the distribution and stability of each metric across different folds. (B) visually presents the performance of classification models on the test set through bar charts, confusion matrices, and ROC curves. (C) the final analysis report, including model parameter selection and key feature sets, and provides a link to download the model for users to further analyze.

## DISCUSSION

### A streamlined but customizable automated machine learning analytics pipeline

The explosive growth of numerical biomedical data presents a challenge in uncovering meaningful insights within the extensive omics data. In recent years, machine learning has emerged as a powerful tool for processing and dissecting big data, making it a popular choice for addressing analytical challenges in biomedical research. Despite providing flexibility and robustness in modeling, the complex algorithms and intricate optimization processes of these model frameworks have deterred many experts in the biomedical field (60). In traditional machine learning analysis workflows, the exploration of numerous classifiers and feature selection algorithms is a common practice. To enhance model performance, various hyperparameters are available for different machine learning algorithms, leading to countless strategy combinations (61). However, this complexity imposes substantial computational burdens and time constraints, serving as a significant barrier for non-expert users attempting to leverage machine learning for data analysis. there exists a critical need for highly automated and expedited analysis tool to optimize the intricate and time-consuming processes of machine learning. However, existing machine learning automation software still faces some challenges: the pipeline designs and method applications are not specifically tailored for addressing biomedical problems, demanding a certain proficiency in programming skills and proving challenging for non-machine learning experts. Some solely concentrate on model selection and parameter optimization, neglecting the pivotal aspect of feature selection, which is essential for high-dimensional data analysis (62).

There is an urgent demand for a straightforward and user-friendly online analysis platform to simplify machine learning analysis and prediction in biomedical field. In response, we have introduced Maler, an automated pipeline designed for online machine learning analysis. Maler selects widely used models in biomedical research, such as support vector machines, random forests, XGBoost, naive Bayes, and others. The Maler analysis pipeline significantly streamlines the steps of machine learning analysis while delivering competitively performing models. For instance, it employs fast and optimal feature ranking and subset search methods, limits the number of features after ranking, and establishes predefined classifier hyperparameters to reduce computational time. These constraints provide researchers with an accessible and user-friendly analysis pipeline, enabling them to generate machine learning analysis results in a short time without the necessity for coding.

### Fast and Scalable Feature Selection Process

Omics datasets often encompass over ten thousand features, yet only a handful of them are crucial for task objectives. Training machine learning models with all features directly can lead to significant computational time consumption and may impact model performance, hindering the discovery of potential biomarkers (63). Therefore, feature selection is an indispensable step to eliminate noise and enhance model computational efficiency. To simplify feature input, retain essential features, and save computational costs, Maler implements a feature ranking and subset selection process.

Among numerous feature selection methods, we ultimately chose ANOVA and MRMR due to their fast and scalable nature as filter-based feature selection methods, offering simplicity and efficiency (64). They exhibit excellent performance in biomedical data. ANOVA, as a univariate feature selection method, rapidly and robustly acquires features correlated with the target variable. On the other hand, MRMR, as a multivariate feature selection method, not only ranks features based on the highest relevance to the target category but also minimizes the mutual correlation between features, eliminating redundancy. Hence, we chose to exclusively provide these methods within Maler.

### The interpretability of signatures remains a challenge

The revival of artificial intelligence presents significant opportunities for the analysis of omics data and research in the biomedical field. However, it also comes with substantial challenges that need to be addressed. The inherent “black-box” nature of machine learning models often makes it difficult for us to interpret and validate biomarkers through computational means (65). This characteristic frequently hinders our ability to biologically explain how models select features for optimal decision-making, limiting their utility in providing insights into potential biological mechanisms and clinical applications.

Researchers have initiated efforts to develop machine learning methods that are biologically interpretable, such as ECMarker (66). However, its current application is limited to classifying cancer phenotypes (staging) based on gene expression, and the classification accuracy still needs to be improved. Consequently, continuous refinement of machine learning algorithms remains essential to establish biologically interpretable models, elucidating the relationship between specific models and input features.

## CONCLUSION

Here, a web server, Maler, is introduced to construct an automated machine learning analysis pipeline. It aims to achieve excellent predictive results by discovering optimal combinations of features and machine learning models. Specifically tailored for researchers in the medical field, Maler server as a user-friendly and powerful solution to help identify and predict relevant features, essential for developing personalized medical approaches and conducting in-depth research on new treatment targets. A key strength of Maler lies in its high degree of automation in machine learning analysis steps, including data preprocessing, feature selection, automatic optimization of machine learning models, and result visualization. This level of automation significantly eases the workflow, particularly for users who may not be proficient in machine learning techniques. Simultaneously, for professional users, Maler provides options to configure hyperparameters, allowing for highly customized analysis workflows. Notably, Maler stands out in expediting biomarker discovery. By employing feature ranking to reduce the dimensionality of high-dimensional omics data, it offers various feature subset search methods based on different models for further feature selection, revealing the most suitable classifier for a given dataset. While demonstrating superior performance compared to similar tools in computational biology through testing with omics data, Maler also exhibits compatibility with any other non-biological data. Finally, Maler utilizes a highly-maintained machine learning library, ensuring easy integration of new implementation algorithms in future versions to maintain the platform’s cutting-edge capabilities and flexibility.

## DATA AVAILABILITY

MALER webserver can be accessed at http://www.inbirg.com/maler/home.

## SUPPLEMENTARY DATA

Supplementary data to this article can be found online at https://github.com/zhaizhaoyu98/maler.

